# Genomic comparison of eight closed genomes of multidrug resistant *Salmonella enterica* strains isolated from broiler farms and processing plants in Trinidad and Tobago

**DOI:** 10.1101/2021.05.20.445003

**Authors:** Meghan Maguire, Anisa S. Khan, Abiodun A. Adesiyun, Karla Georges, Narjol Gonzalez-Escalona

## Abstract

*Salmonella enterica* is an important foodborne pathogen worldwide. We used long and short read sequencing to close genomes of eight multidrug resistant (MDR) *Salmonella enterica* strains, belonging to serovars Infantis (2), Albany, Oranienburg, I 4,[5],12:i:-, Javiana, Schwarzengrund, and Kentucky from broiler chicken farms and processing plants in Trinidad and Tobago. They also belonged to seven different sequence types (STs- 32, 292, 1510, 19, 24, and 96). Seven of the strains demonstrated multi-drug resistance with the presence of at least three AMR genes. Three isolates contained the quinolone resistance gene *qnr*_B19_ in plasmids (CFSAN103840, CFSAN103854, and CFSAN103872). The extended spectrum β-lactamase genes *bla*_CTX-M-65_ (CFSAN103796) and *bla*_TEM-1_ (CFSAN103852) were detected in this study. The genomes closed in this study will be useful for future source tracking and outbreak investigations in Trinidad and Tobago and worldwide.

## Introduction

*Salmonella enterica* is one of the most important foodborne pathogens in the world (González-Escalona et al. 2009). There are more than 325,000 *Salmonella* genomes in NCBI encompassing isolates from many countries (https://www.ncbi.nlm.nih.gov/pathogens/). Nontyphoidal *Salmonella* (NTS) is responsible for approximately 93,757,000 cases of gastroenteritis and 155,000 deaths worldwide each year. Nearly 86% are linked to foodborne transmission (Majowicz et al. 2010). Poultry, including eggs and egg products, is one of the main reservoirs of *Salmonella* and has often been implicated as the source of many outbreaks worldwide (Antunes et al. 2016; Schoeni et al. 1995; Kuehn 2010; Deng et al. 2014; Jackson et al. 2013).

*Salmonella* serovars Enteritidis, Newport, Typhimurium, Javiana, I 4,[5],12:i:-, and Heidelberg have been among the top five serovars associated with human illness over the last 15 years (2006-2016) in the United States ([CSL STYLE ERROR: reference with no printed form.]). *S.* serovar Infantis has shown the highest percent increase in reported cases over the same time frame (+168.5%). While, *S.* serovar Kentucky has become the most commonly detected serovar in chicken (Foley et al. 2011), but is not among the top 20 serovars causing human illness ([CSL STYLE ERROR: reference with no printed form.]).

While most *Salmonella* infections result in gastroenteritis without the need for additional medical intervention, severe, invasive infections can require antimicrobial treatment. Multidrug resistance (MDR) in *Salmonella* is closely monitored in clinical and environmental isolates around the world [National Antimicrobial Resistance Monitoring System (NARMS), European Center for Disease Control and Prevention (ECDC), and European Food Safety Authority (EFSA)]. In the United States, the FDA has issued guidance on the judicious use of antimicrobial agents in food-producing animals (https://www.fda.gov/regulatory-information/search-fda-guidance-documents/cvm-gfi-209-judicious-use-medically-important-antimicrobial-drugs-food-producing-animals) and individual antimicrobial agent restrictions have resulted in decreased prevalence of resistance-conferring genes (https://www.fda.gov/media/108304/download).

Several antimicrobial resistant (AMR) *Salmonella* strains were isolated during a *Salmonella* and AMR surveillance program of poultry production facilities in Trinidad and Tobago during 2016-2019 (Khan, et al, unpublished results). Their genomes were sequenced by short read sequencing and we selected a group of 8 strains belonging to multiple serovars and showing a diverse range of MDR, to completely close their genomes using a combination of long-read and short-read sequencing technologies. The availability of these closed genomes will be useful for future outbreak investigations and for understanding AMR acquisition and transmission in *Salmonella* within the local in Trinidad and Tobago region and worldwide.

## Results and Discussion

An AMR surveillance program of poultry farms and retail outlets in Trinidad and Tobago from 2016 – 2019 produced several *Salmonella* isolates that were confirmed to be *Salmonella* by selective microbiological plating. Initial sequencing efforts identified the sequence type (ST) and serovar for each sample by *in silico* analysis. We selected eight samples from multiple serovars to completely close genomes using a hybrid assembly with Illumina MiSeq short-read and Oxford Nanopore long-read sequencing. The short-read sequencing for these eight samples was performed over three sequencing runs with an average output of 12.6 Gb and estimated average genome coverage of 50 – 150X. The multiplexed nanopore sequencing for eight samples was carried out over two sequencing runs. The output for run 1 was 7.69 Gb (2.33 million reads with a quality score of 9.48 and N50 – 11,835 bp) and for run 2 was 3.71 Gb (694 thousand reads with a quality score of 9.93 and N50 – 12,930 bp). The estimated average genome coverage was 59 - 250X (Table 1).

**Table 1.**
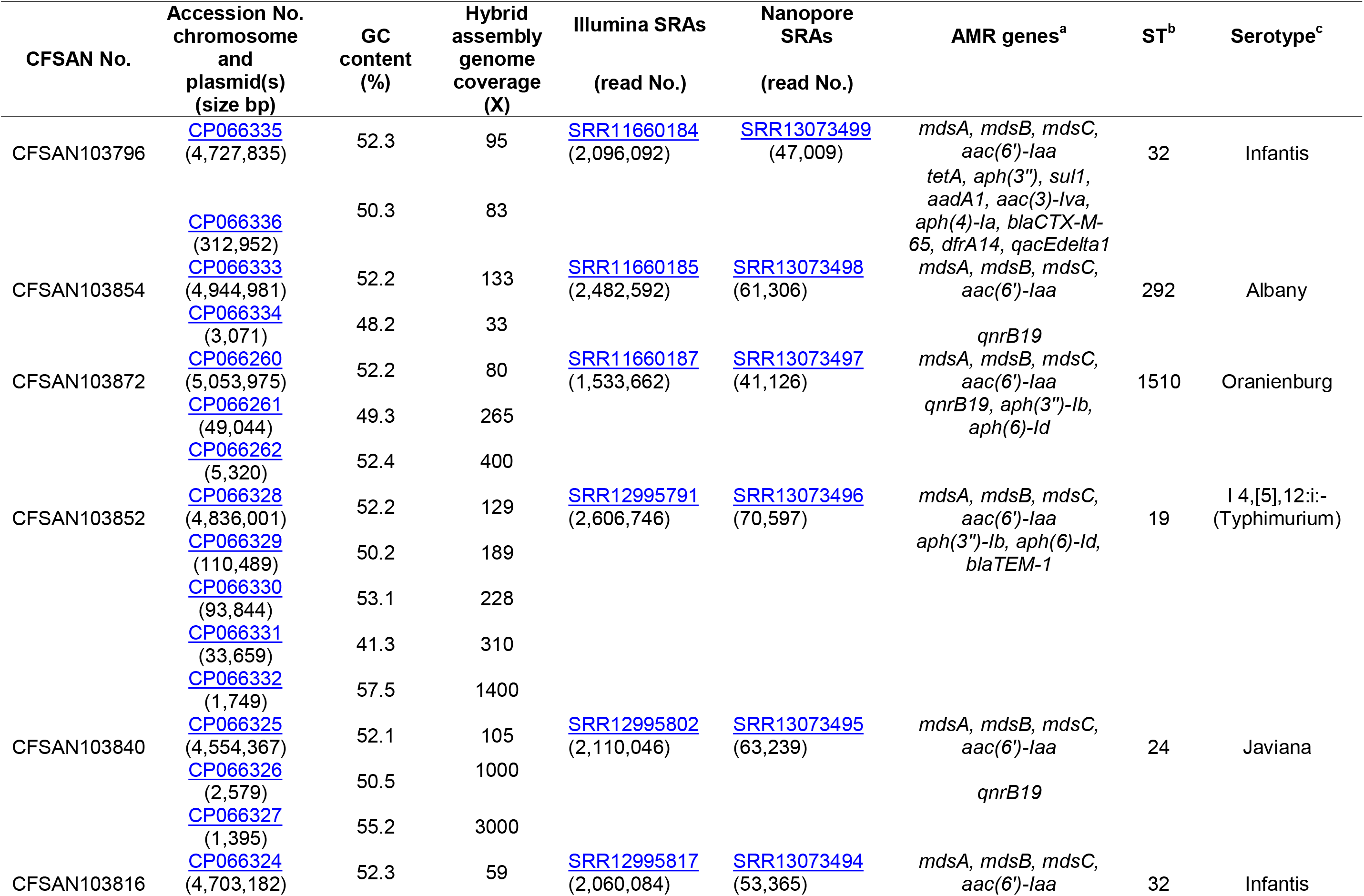
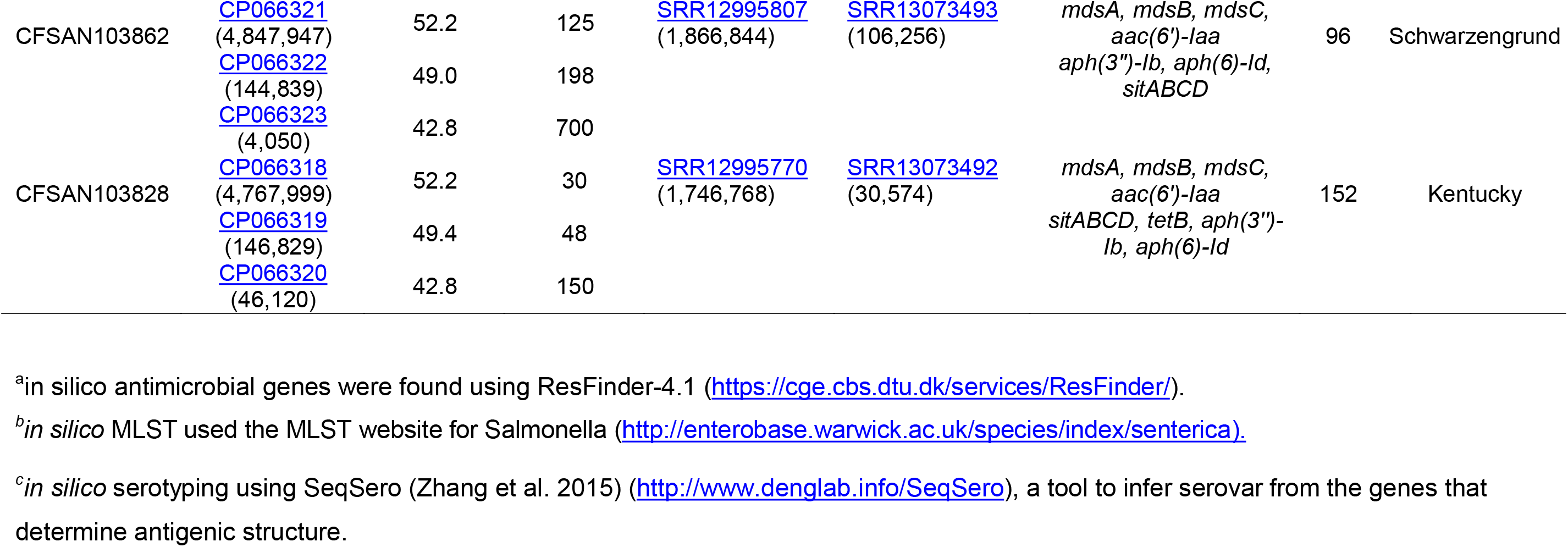
Metadata for the eight *S. enterica* strains isolated from chicken on poultry farms and processing plants reported in this study.

The final closed genomes were generated by a hybrid assembly (Unicycler), which matched in synteny and size to a *de novo* assembly generated using only nanopore reads by the Flye assembler. *In silico* analysis confirmed the ST and serovar for each closed genome observed with the preliminary analysis run using the short-read sequencing (unpublished results) (Table 1). The strains represented seven different STs and serotypes (32 – Infantis, 292 – Albany, 1510 – Oranienburg, 19 - 1,4,[5],12:i:- (Typhimurium), 24 – Javiana, 96 – Schwarzengrund and 152-Kentucky). The genome size of these NTS strains varied from 4.6 Mb (Javiana, CFSAN103840) to 5.1 Mb (Oranienburg, CFSAN103872), highlighting the plasticity of NTS genomes (Table 1). A cgMLST analysis showed that these strains were highly divergent but shared at least 3432 genes out of a total of 4160 genes of the reference *S*. Enteritidis P125109 (NC_011294) strain used for this cgMLST analysis (Figure 1A). A minimum spanning tree of the cgMLST analysis showed that they differed in most loci, >78% (Figure 1B). With the two Infantis (ST32) differing in at least 136 loci, which was expected since Infantis strain CFSAN103816 lacked the MDR plasmid, indicating that these two strains belonged to very different clones circulating in the same country at the same time.

**Figure 1.**
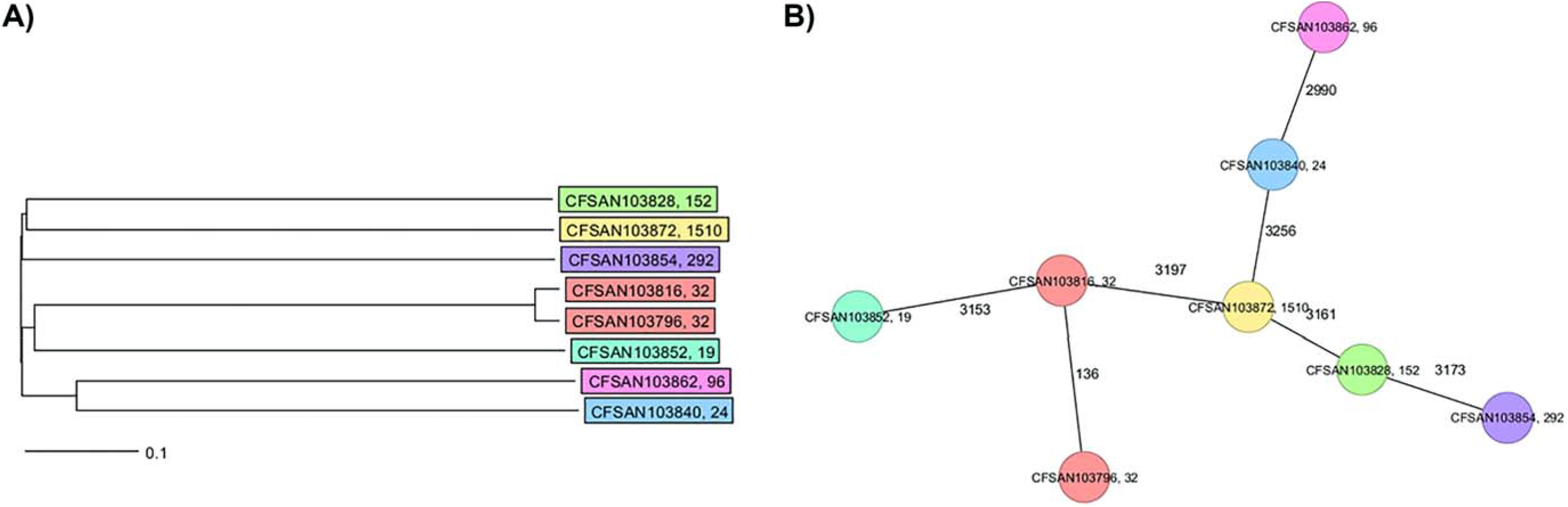
cgMLST analysis of the eight *Salmonella* closed genomes based on 4028 shared genes. A) NJ tree showing the genetic differences among these genomes based on the shared loci. B) Minimum spanning tree allowing to visualize the number of loci differences between these genomes. Strains are color coded according to their ST. The ST is displayed after the name of the strain, separated by a comma. Numbers by lines represent the number of loci differing between strains. The lines are not drawn to scale.

The closed genomes were further analyzed for the presence of AMR genes using the ResFinder database (Table 1) (Bortolaia et al. 2020). All strains were found to contain *mdsA*, *mdsB*, and *mdsC* components of the MdsABC complex in the chromosome. MdsABC confers resistance to multiple toxic compounds, including novobiocin, acriflavine, crystal violet, ethidium bromide, methylene blue, rhodamine 6G, tetraphenylphosphonium bromide, benzalkonium bromide, and SDS and oxidative stress-inducing compounds (diamide, H_2_O_2_, Paraquat), and is necessary for colonization of host cells (Nishino et al. 2006; Song et al. 2015)-. Seven of the strains demonstrated multi-drug resistance with the presence of at least three AMR genes. Aminoglycoside, tetracycline, and sulfonamide resistance genes were detected in seven strains: CFSAN103796 (*aac (6’)*, *tetA*, *aph(3”)*, *sul1*, *aadA1*, *aac(3)-lva*, *aph(4)-la*), CFSAN103854 (*aac(6’)-Iaa*), CFSAN103872 (*aac(6’)-Iaa, aph(3”)-lb*, *aph(6)-ld*), CFSAN103852 (*aac(6’)-Iaa, aph(3”)-lb*, *aph(6)-ld*), CFSAN103816 (*aac(6’)-Iaa*), CFSAN103862 (*aac(6’)-Iaa, aph(3”)-lb*, *aph(6)-ld*), and CFSAN103828 (*aac(6’)-Iaa, aph(3”)-lb*, *aph(6)-ld*, *tetB*) (Table 1). In clinical *Salmonella* isolates, combined resistance to ampicillin, chloramphenicol, streptomycin, sulfonamides and tetracycline (ACSSuT) decreased from 8.2% in 1996 to 3.6% in 2009 (Medalla et al. 2013), but other medically relevant antibiotics have become a recent focus of study. Synthetically derived fluoroquinolones were introduced to combat ACSSuT resistance. Three isolates contained the quinolone resistance gene *qnr*_B19_ in plasmids (CFSAN103840, CFSAN103854, and (CFSAN103872) of different sizes. Plasmid-mediated quinolone resistance was not identified in retail meat products until 2017 (Sjölund-Karlsson et al. 2010; Tyson et al. 2017). The presence of this small plasmids carrying *qnr*_B19_ gene is worrisome since they have been identified in other countries in Latin America such as Peru, Bolivia, Colombia, and Argentina and appears to be highly mobile among different microorganisms (Tran et al. 2012).

β-lactams, including ampicillin and cephalosporin, are another treatment of invasive salmonellosis, particularly in children. Ceftriaxone resistance increased in the US from 10% in 2002 to 38% in 2010 in chicken, prompting prohibition of cephalosporin by the FDA in 2012. The extended spectrum β-lactamase genes *bla*_CTX-M-65_ (CFSAN103796) and *bla*_TEM-1_ (CFSAN103852) were detected in this study. Brown, et al (2018) described the presence of *bla*_CTX-M-65-_containing *Salmonella* isolates in the US that were grouped in the same phylogenic clade as MDR Peruvian isolates (Brown et al. 2018), illustrating the intercontinental spread of this AMR gene. We have additionally detected the presence of the *mcr-9* gene in another strain isolated with this study (Maguire et al. 2021). The *mcr-9* gene variant was described in 2019 (Carroll et al. 2019) and appears not to confer colistin resistance in *Salmonella* and *E. coli* (Tyson et al. 2020).

## Conclusion

MDR NTS presence in poultry environments remains a significant global challenge for public health. With the increasing reports of newer variants of AMR genes that confer resistance to diverse types of antibiotics, the need to enact active AMR gene surveillance studies is paramount. With these types of studies, we can not only detect the presence and spread of important AMR genes in poultry, but also identify the strains carrying these AMR genes and their location (whether in the chromosome or plasmids). Here we closed eight genomes of several MDR NTS, determine their ST, serotype, and their associated AMR gene profiles, which will be useful for future source tracking and outbreak investigations in Trinidad and Tobago and worldwide.

## Materials and Methods

### Bacterial strains and media

Cloacal swabs, drag swabs, whole chicken carcases, and neck skin samples were collected from retail outlets, processing plants, and farms in Trinidad and Tobago during an AMR surveillance program from 2016 – 2019. The sample was pre-enriched in buffered peptone water (Oxoid, Ltd., Hampshire, UK) for 18 to 24 h at 37°C, then selectively enriched in tetrathionate broth (Oxoid), and thereafter incubated for 18 to 24 h at 37°C. The sample enriched in selective broth was then sub-cultured onto xylose lysine Tergitol 4 (Oxoid) and incubated aerobically at 37°C for 18 to 24 h. Suspected Salmonella colonies that displayed characteristic colonies on the selective agar plate were then purified on blood agar plates (Oxoid) and incubated at 37°C for 18 to 24 h. Pure cultures were subjected to a panel of biochemical tests that included triple sugar iron agar, lysine iron agar, urea, citrate, methyl red, sulfide-indole-motility medium, and o-nitrophenyl-b-D-galactopyranoside (Oxoid) (Andrews 1992). Biochemically confirmed isolates were then subjected to serological typing by using Salmonella polyvalent antiserum (A-I and Vi, Difco, Detroit, MI). Complete confirmation and serotyping of Salmonella isolates were performed by using the phase reversal technique, and the results interpreted according to the Kauffman-White scheme (Grimont & Weill 2007) at the Public Health Laboratory, Ministry of Health, St. Michael, Barbados. The single isolate strains were grown overnight in tryptic soy broth (TSB) medium at 35°C.

### Nucleic acid extraction

DNA was extracted using the Maxwell RSC cultured cells DNA kit with a Maxwell RSC instrument (Promega, Madison, WI) following the manufacturer’s protocols for Gram-negative bacteria with additional RNase treatment. DNA concentration was determined by Qubit 4 Fluorometer (Invitrogen, Carlsbad, CA) according to manufacturer’s instructions.

### Illumina MiSeq short-read sequencing

The short reads whole genome sequence for this strain was generated by Illumina MiSeq sequencing with the MiSeq V3 kit using 2 × 250 bp paired-end chemistry, (Illumina, San Diego, CA) according to manufacturer’s instructions. The libraries were constructed using 100 ng of genomic DNA using the Illumina® DNA Prep (M) Tagmentation (Illumina), according to manufacturer’s instructions. Reads were trimmed with Trimmomatic v0.36 (Bolger et al. 2014).

### Oxford Nanopore long-read sequencing

The long reads for each strain were generated through GridION sequencing (Nanopore, Oxford, UK). The sequencing library was prepared using the rapid barcoding sequencing kit (SQK-RBK004). Each library was run in a FLO-MIN106 (R9.4.1) flow cell, according to the manufacturer’s instructions, for 48 hours. Default parameters were used for all software unless otherwise specified. The run was base called live with default settings (MinKNOW Core v3.6.5, Bream v4.3.16, and Guppy v3.2.10). Reads < 4000 bp and quality scores of <7 were discarded for downstream analysis for an

### Contig assembly and annotation

The final complete genome (chromosome and plasmid(s), when present) for each strain was obtained using a previously described pipeline (Gonzalez-Escalona et al. 2018) with modifications. The final genomes were achieved by a hybrid assembly generated using both nanopore and MiSeq data with Unicycler v0.4.8 (Wick et al. 2017). A second *de novo* genome assembly was obtained with Flye v2.8 (Kolmogorov et al. 2019) using nanopore data. Both assemblies (Flye and Unicycler) for each strain were aligned with Mauve v2.4.0 (Darling et al. 2004). If the two aligned assemblies agreed in synteny and size, then the Unicycler hybrid assembly was used as the final assembly (i.e. complete genome). Unicycler assembled the chromosomes and plasmids as circular closed and oriented the chromosomes to start at the *dnaA* gene. The genomes were annotated using the NCBI Prokaryotic Genomes Automatic Annotation Pipeline (PGAAP, http://www.ncbi.nlm.nih.gov/genome/annotation_prok) (Tatusova et al. 2016). *In silico* analysis was used to determine the multi-locus sequence type (MLST) with the MLST website for *Salmonella* (http://enterobase.warwick.ac.uk/species/index/senterica), serotype using SeqSero (Zhang et al. 2015) (http://www.denglab.info/Seqero) which infers the serovar from antigenic structure, and antimicrobial genes were found using ResFinder-4.1 (https://cge.cbs.dtu.dk/services/ResFinder/) (Bortolaia et al. 2020).

### Genomic data analysis

To analyze the relationships among the eight NTS closed genomes we performed a core genome MLST (cgMLST) using Ridom Seqsphere+ v6.0.2 (Ridom GmbH, Germany) as previously described (Toro et al. 2016). The genome of *S*. Enteritidis P125109 (NC_011294) was used as reference genome (4,160 genes) (Thomson et al. 2008). The *S*. Enteritidis genome from strain EC20121175 (CP007269.2) was used for comparison with the reference genome to establish a list of core and accessory genome genes. Genes that are repeated in more than one copy in any of the two genomes were removed from the analysis as failed genes. A task template was created that contains both core and accessory genes for this reference SE strain for any future testing. Each individual locus (core or accessory genes) from strain P125109 was assigned allele number 1. The assemblies for each individual NTS closed genome in this study were queried against the task template and if the locus was found and was different from the reference genome or any other queried genome already in the database, a new number was assigned to that locus and so on. After eliminating any loci that were missing from the genome of any strain used in our analyses, we performed the cgMLST analysis. These remaining loci were considered the core genome shared by the analyzed strains. We used Nei’s DNA distance method (Nei et al. 1983) for calculating the matrix of genetic distance, taking only the number of same/different alleles in the core genes into consideration. A Neighbor-Joining (NJ) tree using the appropriate genetic distances was built after the cgMLST analysis.

### Data deposition

The complete genome sequences of the eight *S. enterica* strains have been deposited at GenBank under the accession listed in Table 1.

## REFERENCES

Andrews W. 1992. Manual of food quality control. 4. Rev. 1. Microbiological analysis. Food and Drug Administration. FAO Food Nutr. Pap. 14:1–338.

Antunes P, Mourão J, Campos J, Peixe L. 2016. Salmonellosis: the role of poultry meat. Clin. Microbiol. Infect. 22:110–121. doi: 10.1016/j.cmi.2015.12.004.

Bolger AM, Lohse M, Usadel B. 2014. Trimmomatic: a flexible trimmer for Illumina sequence data. Bioinformatics. 30:2114–2120. doi: 10.1093/bioinformatics/btu170.

Bortolaia V et al. 2020. ResFinder 4.0 for predictions of phenotypes from genotypes. J. Antimicrob. Chemother. 75:3491–3500. doi: 10.1093/jac/dkaa345.

Brown AC et al. 2018. CTX-M-65 extended-spectrum β-lactamase–producing *Salmonella enterica* serotype Infantis, United States. Emerg. Infect. Dis. 24:2284–2291. doi: 10.3201/eid2412.180500.

Carroll LM et al. 2019. Identification of novel mobilized colistin resistance gene *mcr-9* in a multidrug-resistant, colistin-susceptible *Salmonella enterica* serotype Typhimurium isolate. MBio. 10:e00853–19. doi: 10.1128/mBio.00853-19.

Darling ACE, Mau B, Blattner FR, Perna NT. 2004. Mauve: Multiple alignment of conserved genomic sequence with rearrangements. Genome Res. 14:1394–1403. doi: 10.1101/gr.2289704.

Deng X et al. 2014. Genomic epidemiology of *Salmonella enterica* serotype Enteritidis based on population structure of prevalent lineages. Emerg. Infect. Dis. 20:1481–1489. doi: 10.3201/eid2009.131095.

Foley SL et al. 2011. Population dynamics of *Salmonella enterica* serotypes in commercial egg and poultry production. Appl. Environ. Microbiol. 77:4273–4279. doi: 10.1128/AEM.00598-11.

González-Escalona N et al. 2009. Detection of live *Salmonella* sp. cells in produce by a TaqMan-based quantitative reverse transcriptase real-time PCR targeting *invA* mRNA. Appl. Environ. Microbiol. 75:3714–3720. doi: 10.1128/AEM.02686-08.

Gonzalez-Escalona N, Haendiges J, Miller JD, Sharma SK. 2018. Closed genome sequences of two *Clostridium botulinum* strains obtained by nanopore sequencing. Microbiol. Resour. Announc. 7:e01075–18. doi: 10.1128/MRA.01075-18.

Grimont PAD, Weill F-X. 2007. Antigenic formulae of the *Salmonella* serovars. WHO Collaborating Centre for Reference and Research on Salmonella, Paris.

Jackson BR, Griffin PM, Cole D, Walsh KA, Chai SJ. 2013. Outbreak-associated *Salmonella enterica* serotypes and food Commodities, United States, 1998-2008. Emerg. Infect. Dis. 19:1239–1244. doi: 10.3201/eid1908.121511.

Kolmogorov M, Yuan J, Lin Y, Pevzner PA. 2019. Assembly of long, error-prone reads using repeat graphs. Nat. Biotechnol. 37:540–546. doi: 10.1038/s41587-019-0072-8.

Kuehn BM. 2010. *Salmonella* cases traced to egg producers. JAMA. 304:1316. doi: 10.1001/jama.2010.1330.

Maguire M, Khan AS, Adesiyun AA, Georges K, Gonzalez-Escalona N. 2021. Closed genome sequence of a *Salmonella enterica* serotype Senftenberg strain carrying the *mcr-9* gene isolated from broken chicken eggshells in Trinidad and Tobago. Microbiol. Resour. Announc. 10:e01465–20. doi: 10.1128/MRA.01465-20.

Majowicz SE et al. 2010. The global burden of nontyphoidal *Salmonella* gastroenteritis. Clin. Infect. Dis. 50:882–889. doi: 10.1086/650733.

Medalla F et al. 2013. Increase in Resistance to Ceftriaxone and nonsusceptibility to Ciprofloxacin and decrease in multidrug resistance among *Salmonella* strains, United States, 1996-2009. Foodborne Pathog. Dis. 10:302–309. doi: 10.1089/fpd.2012.1336.

National Enteric Disease Surveillance: *Salmonella* Annual Report, 2016. 87.

Nei M, Tajima F, Tateno Y. 1983. Accuracy of estimated phylogenetic trees from molecular data. II. Gene frequency data. J.Mol.Evol. 19:153–170.

Nishino K, Latifi T, Groisman EA. 2006. Virulence and drug resistance roles of multidrug efflux systems of *Salmonella enterica* serovar Typhimurium. Mol. Microbiol. 59:126–141. doi: https://doi.org/10.1111/j.1365-2958.2005.04940.x.

Schoeni JL, Glass KA, McDermott JL, Wong ACL. 1995. Growth and penetration of *Salmonella* enteritidis, *Salmonella* heidelberg and *Salmonella* typhimurium in eggs. Int. J. Food Microbiol. 24:385–396. doi: 10.1016/0168-1605(94)00042-5.

Sjölund-Karlsson M et al. 2010. Plasmid-mediated quinolone resistance among non-Typhi *Salmonella enterica* isolates, USA. Emerg. Infect. Dis. 16:1789–1791. doi: 10.3201/eid1611.100464.

Song S et al. 2015. MdsABC-mediated pathway for pathogenicity in *Salmonella enterica* serovar Typhimurium. Infect. Immun. 83:4266–4276. doi: 10.1128/IAI.00653-15.

Tatusova T et al. 2016. NCBI prokaryotic genome annotation pipeline. Nucleic Acids Res. 44:6614–6624. doi: 10.1093/nar/gkw569.

Thomson NR et al. 2008. Comparative genome analysis of *Salmonella* Enteritidis PT4 and *Salmonella* Gallinarum 287/91 provides insights into evolutionary and host adaptation pathways. Genome Res. 18:1624–1637. doi: 10.1101/gr.077404.108.

Toro M et al. 2016. Whole-genome sequencing analysis of *Salmonella enterica* serovar Enteritidis isolates in Chile provides insights into possible transmission between gulls, poultry, and humans. Appl.Environ.Microbiol. 82:6223–6232. doi: 10.1128/AEM.01760-16.

Tran T et al. 2012. Small plasmids harboring *qnrB19*: a model for plasmid evolution mediated by site-specific recombination at *oriT* and *Xer* sites. Antimicrob. Agents Chemother. 56:1821–1827. doi: 10.1128/AAC.06036-11.

Tyson GH et al. 2017. Identification of plasmid-mediated quinolone resistance in *Salmonella* isolated from swine ceca and retail pork chops in the United States. Antimicrob. Agents Chemother. 61. doi: 10.1128/AAC.01318-17.

Tyson GH et al. 2020. The *mcr-9* gene of *Salmonella* and *Escherichia coli* is not associated with colistin resistance in the United States. Antimicrob. Agents Chemother. 64:e00573–20. doi: 10.1128/AAC.00573-20.

Wick RR, Judd LM, Gorrie CL, Holt KE. 2017. Unicycler: Resolving bacterial genome assemblies from short and long sequencing reads. PLOS Comput. Biol. 13:e1005595. doi: 10.1371/journal.pcbi.1005595.

Zhang S et al. 2015. *Salmonella* serotype determination utilizing high-throughput genome sequencing data J. Clin. Microbiol. 53:1685. doi: 10.1128/JCM.00323-15.

